# Decline of intrinsic cerebrospinal fluid outflow in healthy humans with age detected by non-contrast spin-labeling MRI

**DOI:** 10.1101/2022.07.14.500033

**Authors:** Vadim Malis, Won C. Bae, Asako Yamamoto, Linda K. McEvoy, Marin A. McDonald, Mitsue Miyazaki

**Author notes:** Corresponding Author: Mitsue Miyazaki, Ph D, Department of Radiology, University of California San Diego 9472 Health Science Drive, La Jolla, CA 92093 USA.

## Abstract

**Background:** Clearance of cerebrospinal fluid (CSF) is important for the removal of toxins from the brain, with implication for neurodegenerative diseases. Imaging evaluation of CSF outflow in humans has been limited, relying on injections of contrast agents. Objective of this study was to introduce a novel spin-labeling magnetic resonance imaging (MRI) technique to detect and quantify the movement of endogenously tagged CSF without administration of tracer or contrast media, and use the technique to evaluate CSF outflow in normal human subjects with varying ages.

**Methods:** This study was performed on a clinical 3-Tesla MRI scanner in healthy subjects (10 males and 6 females; mean age, 47.6 ± 18.9 years; range, 19-71 years) with informed consent. Our non-contrast spin-labeling MRI technique applies a tag pulse on the brain hemisphere, including subarachnoid space, dura mater, brain parenchyma, and images the outflow of the tagged CSF into the superior sagittal sinus. We obtained 3-dimensional images in real time, which was analyzed to determine tagged-signal changes in different regions of the brain involved in CSF outflow or clearance. Additionally, the signal changes over time were fit to a signal curve to determine quantitative flow metrics such as relative CSF flow and volume. These were correlated against subject age to determine aging effects.

**Results:** We observed the signal of the tagged CSF moving from the subarachnoid space to the dura mater and parasagittal dura, and finally draining into the superior sagittal sinus. In addition, there was strong evidence of a direct pathway by which tagged CSF flows directly from the subarachnoid space to the superior sagittal sinus, via the lateral wall of superior sagittal sinus. Furthermore, quantitative CSF outflow metrics were shown to decrease significantly with age.

**Conclusions:** We demonstrated a novel non-invasive MRI technique to evaluate CSF clearance in humans. In this study, we identified possible two CSF clearance pathways, and determined normative values and decline of CSF flow metrics in normal ages. Our work provides a new opportunity to better understand the relationships of these CSF clearance pathways in ages, which may be a significant factor in the age-related prevalence of neurodegenerative diseases.

**Funding:** This study was supported by the National Institutes of Health grants: RF1AG076692 (MM) and R01HL154092 (MM); and made possible by a grant from Canon Medical Systems. Corp., Japan.

**Clinical trial:** Not applicable.

## INTRODUCTION

The investigation of the pathways of glymphatic fluid clearance has been gaining interest due to the potential association between dysfunction of brain waste clearance and neurodegenerative diseases such as Alzheimer’s disease (*Nedergaard, 2013*). The concept of glymphatic fluid clearance in the brain was pioneered by Nedergaard, who identified a system by which soluble proteins and metabolites are eliminated from the central nervous system via cerebrospinal fluid (CSF) and interstitial fluid exchange in the paravascular space (*Iliff et al., 2012; 2013*). Our understanding of the importance of this system continues to expand, predominantly from studies using either ionizing radiation or invasive contrast injection in rodents (*Nedergaard, 2013; Iliff et al., 2012; 2013; Benveniste et al., 2017; Jessen et al., 2015; Kress et al., 2014*). However, rodent models may not fully recapitulate the human glymphatic system, necessitating study of CSF flow in humans (*Ringstad et al., 2020*).

The classic glymphatic drainage pathway has been considered to flow from the subarachnoid space (SAS) to the superior sagittal sinus (SSS), via arachnoid granulations (AGs) (*Pollay 2010; Sakka et al., 2011*) (see Fig 1). However, studies using invasive methodologies in animals, have identified a second pathway involving dural lymphatic vessels (*Aspelund et al., 2015; Louveau et al., 2015; Da Mesquita et al., 2018*). Gadolinium-based contrast agent (GBCA) magnetic resonance imaging (MRI) studies have revealed dural lymphatic vessels in human and nonhuman primates as well (*Absinta et al., 2017*). It is not yet known whether other glymphatic drainage pathways may also exist in human.

**Fig. 1.**
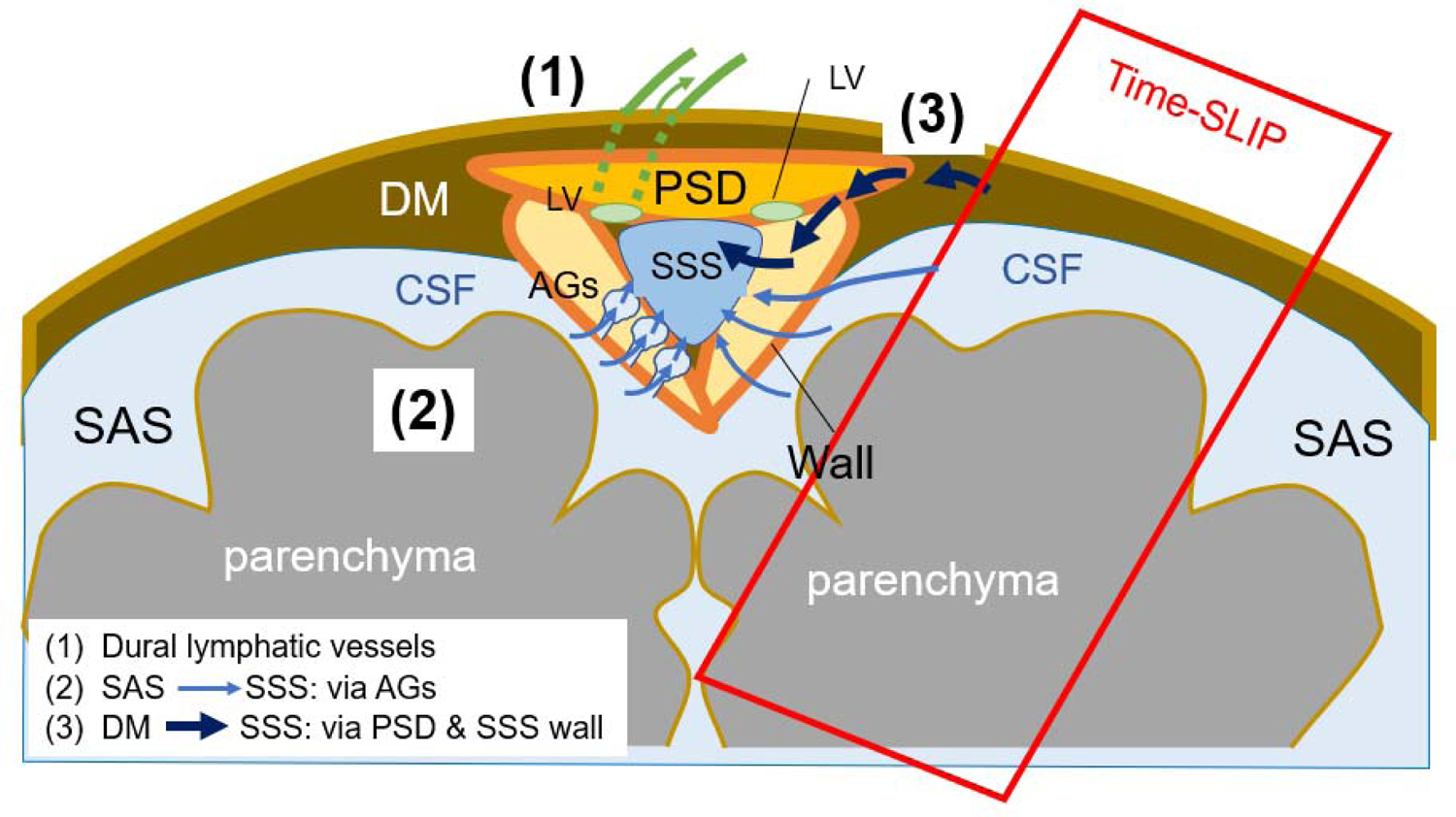
Drawing of possible egress pathways of CSF. (1) Pathway of egress through dural lymphatic vessels, observed using tracer studies, including gadolinium-based contrast agent. (2) Classic theory of CSF egress from subarachnoid space (SAS) to the superior sagittal sinus (SSS), via arachnoid granulations (AGs). (3) Pathway of CSF from the dura mater (DM) to parasagittal dura (PSD), and then to SSS, observed using our tagged CSF method.

Recently, the use of GBCA via intrathecal injection in humans revealed uptake and washout of the tracer at the meninges as evidence of glymphatic clearance, albeit over an extended, 48-hour period^7^. These GBCA studies provide valuable evidence of dural lymphatic vessels and their connection to parasagittal dura (PSD) and arachnoid granulations (AG) in humans. However, they have not assessed the intrinsic outflow of the glymphatic fluid itself or quantified outflow metrics. Detailed reports on the direction of fluid movement, real-time visualization of glymphatic fluid movement, and quantification of CSF efflux in humans are needed to better understand the CSF flow pathways in humans.

The present study was therefore aimed to elucidate the egress pathways of intrinsic CSF outflow using a novel MRI method that does not require tracer or contrast injections, and to quantify measures of outflow. Because glymphatic function has been shown to change with age, we also examined outflow metrics in younger and older adults as proof of concept and to investigate potential age-related changes in CSF outflow tracts. In this article, we discuss the egress pathways of fluid outflow from the brain, using the term CSF to encompass interstitial fluid (ISF)-CSF fluid, glymphatic fluid, or brain fluid throughout the article.

## MATERIALS AND METHODS

### Experimental design

The study was approved by the Institutional Review Board of University of California, San Diego (200335). Written informed consent was obtained from all consecutive study participants. The study design was observational: Randomization of participants was not required, and we did not perform a prior sample size calculation since this was the first study of its kind.

Our consecutive study participants consisted of 16 healthy adults (10 males and 6 females; age range 19-71 years; mean age, 49.8 ± 13.8 years), each imaged in a supine position. None of the participants had any known neurodegenerative disease, pulmonary disease, or cardiovascular disease. Table 1 shows the participant demographics. Participants were further divided into two groups based on aged: younger (19 to 59 years, n=8) and older (60 years and older, n=8) groups, to better assess age differences. All participants underwent two-dimensional (2D) T2-weighted FLAIR, three-dimensional (3D) T2-weighted centric *k*_y_–*k*_z_ single shot fast spin echo (cSSFSE), and four-dimensional (4D) fluid spin-labeling MRI using 3D cSSFSE with various inversion time (TI) periods. All scans were acquired at mid-day to control for circadian effects.

**Table 1.**
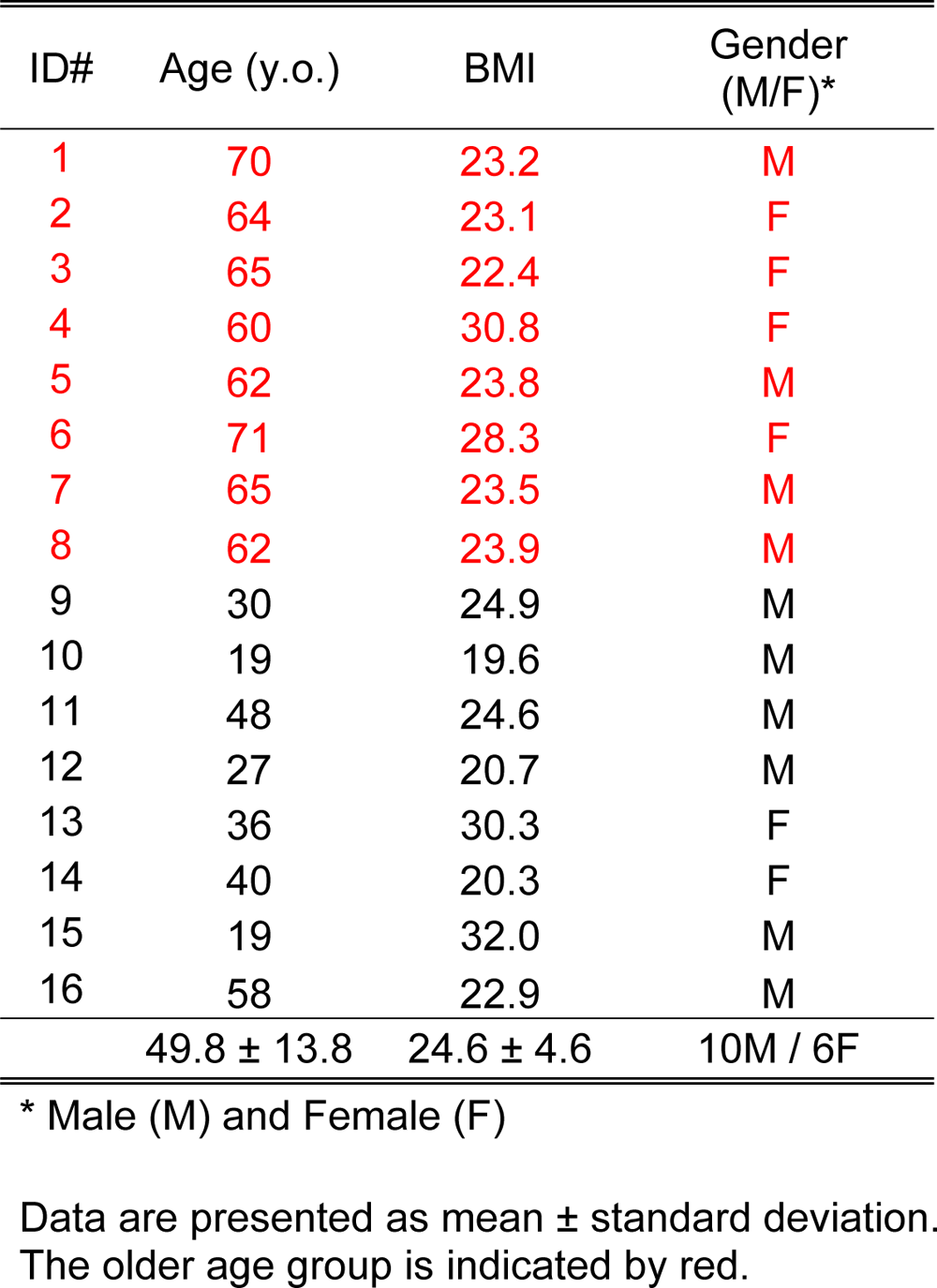
Subject Demographics

### MR protocol

All MRI examinations were performed on a clinical 3-T MRI scanner (Vantage Galan, Canon Medical Systems Corp., Japan), equipped with a 32-ch head coil.

1. T2-weighted 2D fluid attenuated inversion recovery (FLAIR), repetition time (TR)=11000 ms, effective echo time (TE_eff_)=92 ms, echo train spacing (ETS)=11.5 ms, inversion time (TI)=2900 ms, flip angle/refocusing angle = 90/170°, 1 average, 32 slices, 2 mm thick, matrix=384 × 364 (768 × 738 after interpolation), FOV = 22 × 22 cm, parallel imaging factor of 2.4, and acquisition time of 4 min 2 s.
2. T2-weighted 3D cSSFSE parameters are TR=4200 ms, TE_eff_ = 121 ms, ETS = 6.5 ms, number of shots = 1, flip/flop = 90/110°, spectral attenuated inversion recovery (SPAIR) fat suppression, matrix size = 432 × 432, (864 × 864 after interpolation), FOV = 22 × 22 cm, 100 slices, 0.6 mm thick, near isovoxel resolution of 0.3 mm × 0.3 mm × 0.3 mm after interpolation, parallel imaging factor of 3, and acquisition time of 3 min and 26 s. Denoising deep learning reconstruction (dDLR) was applied (*Kidoh et al., 2019*).
3. Ungated 4D CSF spin-labeling MRI parameters with 3D cSSFSE readout are TR=5400 ms, TE_eff_ = 30 ms, ETS = 5.0 ms, number of shots = 1, flip/refocusing angle = 90/150°, SPAIR fat suppression, matrix size =368 × 368 (736 × 736 after interpolation), FOV=25 × 25 cm, parallel imaging factor of 3, 20 slices, 1 mm thick (0.5 mm after interpolation), tag-on and tag-off acquisition time of about 1 min and 48 s. Images were acquired for the inversion time (TI) of 500, 750, 1000, 1250, 1500, 2000, and 3000 ms. Temporal resolution of acquired data points between the TI periods was varied. The higher temporal resolution around 1000 ms was adopted since the preliminary scans showed it to be the TI where the peak flow occurs. Table 2 shows the summary of MR imaging acquisition parameters.

**Table 2.** MR Imaging Acquisition Parameters

| Parameter | Units | T2 FLAIR | 3D cSSFSE | 3D Spin-Labeling |
| --- | --- | --- | --- | --- |
| Field of view | [mm] | 220 × 220 × 64 | 220 × 220 × 60 | 250 × 250 × 20 |
| Image matrix |  | 384 × 364 × 32 | 432 × 432 × 100 | 368 × 368 × 20 |
| No. of sections |  | 32 | NA | NA |
| Slice thickness | [mm] | 2 | 0.6 | 1 |
| Acquired resolution:<br>(PE) × (RO) × (SE) | [mm] | 0.57 × 0.60 × 2.0 | 0.51 × 0.51 × 0.6 | 0.68 × 0.68 × 1.0 |
| Interpolated resolution:<br>(PE) × (RO) × (SE) | [mm] | 0.29 × 0.3 × 1.0 | 0.25 × 0.25 × 0.3 | 0.34 × 0.34 × 0.5 |
| Repetition time | [ms] | 11000 | 4200 | 5400 |
| Effective echo time | [ms] | 92 | 121 | 30 |
| Echo train spacing | [ms] | 11.5 | 6.5 | 5 |
| Flip/fop angles | [degrees] | 90/170 | 90/110 | 90/150 |
| Averages |  | 1 | 1 | 1 |
| Fat suppression (SPAIR) |  | off | on | on |
| Parallel reduction factor |  | 2.4 | 3 | 3 |
| Acquisition time |  | 4 min 2 sec | 3 min 26 sec | 1 min 48 sec<br>(for each TI) |

### 4D spin-labeling MRI theory

Fig. 3A shows the pulse sequence of alternated tag-on and tag-off slice encoding acquisitions with their magnetization and Fig. 3B shows post-processing of tag-on and tag-off in 3D manner, and their subtraction. An adiabatic spatially selective inversion recovery pulse, time-spatial labeling inversion pulse (Time-SLIP), is used in the tag-on acquisition. A tag-on and tag-off alternate acquisition and subtraction technique was used with centric *k*_y_–*k*_z_ k-space ordering 3D SSFSE (cSSFSE). The tag-on sequence consists of a non-selective inversion recovery (non-sel-IR) pulse and a spatial-selective IR (sel-IR) pulse as a labeling (tag) pulse (Fig. 3A left). First, the non-sel IR pulse inverts all signals in the field of view (FOV) from the initial longitudinal magnetization (+*M*_z_) to (– *M*_z_). Second, immediately after the non-sel-IR pulse, the sel-IR pulse is applied to invert back only the magnetized tissue and fluid in the region of interest (ROI). Thus, the longitudinal magnetization in the tagged region is restored to +*M*_z_, whereas the magnetization elsewhere remains at –*M*_z_ and follows an exponential T_1_ relaxation curve, as indicated in Fig. 3A left. The tag-off sequence consists of only non-sel IR pulse, as shown in Fig. 3A right. Both tag-on and tag-off acquisitions were alternately collected, and their subtraction provides flow-out signals from the tag pulse (*Miyazaki et al., 2008; 2011; 2012; 2015; Wheaton et al., 2012*). Both tag-on and tag-off acquisitions have the non-sel IR pulse and thus the background signals with exponential T1 relaxation magnetization returns are the same in both images; therefore, background signals are canceled out and only the tagged signal that moves out of the Time-SLIP pulse is observed. For 3D volume data, this tag-on and tag-off acquisition was repeated in every slice encoding in TR. After this preparation, marked CSF signals flow out from the tag pulse and move out during the TI period (between the tag and read-out).

### Experimental studies: visualization of CSF and effect of aging on outflow metrics

We acquired all 3 protocols described in the MR Protocol section (as well as Table 2.) on all 16 participants. The 3D coronal imaging slab was placed at the top of the head and an oblique tag pulse was applied alongside the superior sagittal sinus (SSS), as shown in Supplementary 1 (S1). The detailed anatomies of dura mater, PSD, SSS wall, AGs, as well as SAS were realized using coronal images of T2-weighted FLAIR and T2-weighted 3D cSSFSE with distinct intrinsic contrast. Our ungated 4D CSF spin-labeling MRI technique was previously optimized to acquire TI of 500, 750, 1000, 1250, 1500, 2000, and 3000 ms, a total of 7 acquisitions (each TI with 1 min 48 sec.) to have higher temporal resolution of TI around 1000 ms, where a peak height (PH) was observed. As shown in Fig. 1 (3), our hypothesis is that tagged CSF in dura mater travels to parasagittal dura (PSD) and superior sagittal sinus (SSS) wall, where arachnoid granulations (AGs) are located, and then flows out into the SSS. We have studied both simple subtraction of tag-on and tag-off images as well as signal increase ratio (SIR), after registration of both images. The detailed calculation of SIR is discussed in the Data analysis and post-processing.

Along with the main experiment, we investigated the reproducibility of our methodology. We consecutively acquired tagged CSF outflow within a single scanning session in two participants to avoid differences in CSF flow related to circadian effects, or differences in prior physical activity or sleep.

### Data analysis and post-processing

We measured four ROIs of PSD, SSS wall, SSS, as well as the entire SSS region containing PSD, SSS lateral walls, and SSS. Regions were manually selected for each participant on the tag-on image (TI = 500 ms) away from the tagged region by an MR physicist (MM) and a medical doctor (XZ) together. We carefully selected the 4 ROIs from images of TI of 500 ms together with T2-weighted FLAIR and T2-weighted 3D cSSFSE because the image contrast of TI of 500 ms provides well-defined regions of PSD and SSS walls in bright signal with dark SSS signal. For both volumes (tag-on and tag-off) voxels with intensity values outside the range [MEAN ± 2SD] were excluded from further analysis. We performed this method to eliminate voxels that may contain signals of pulsatile blood flow between the tag-on and tag-off images and between TI periods. This ensures that our tagged MRI signal is indeed from the CSF and not the blood.

Normal distribution of intensities in tag-on and tag-off images is expected and the range of [MEAN ± 2SD] contains 95% of the voxels in the selected region of interest (*Huber 2018*). To determine the percent signal increase ratio (SIR) from outflow, tag-off images were subtracted from tag-on images, and the absolute value of the difference was then divided by tag-off images, as shown in equation [1].

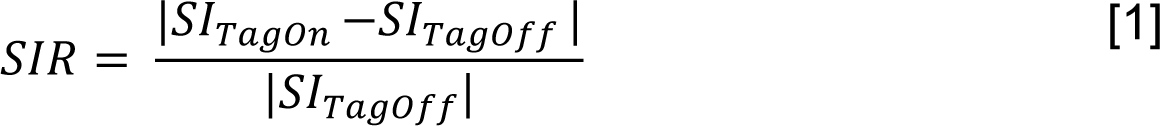

This was performed for each TI, resulting in a time-varying signal increase at each voxel. The ROI averaged values (y) were then used to perform least-squares fit for the TI-dependent gamma variate function (*Chan et al., 2004*) given by equation [2]:

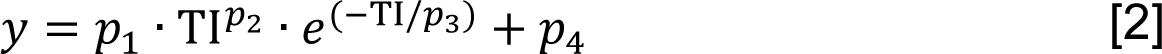

The starting points *p^0^_i_* and range of the parameters used for fit were:

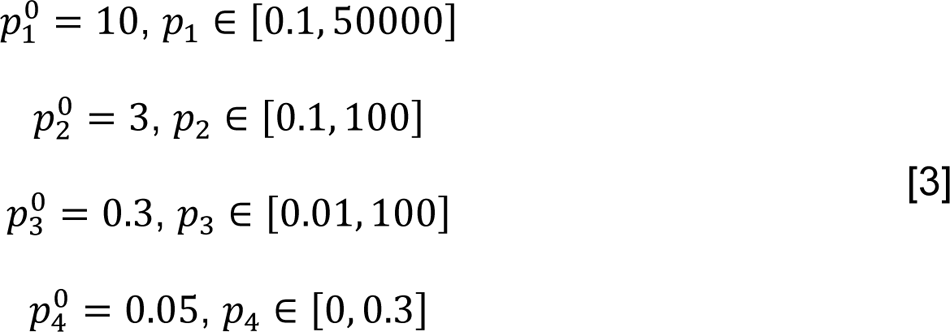

Fitting results produced estimates for five outflow metrics (*Copen et al., 2011*) including: peak height (PH), time-to-peak (TTP), mean transit time (MTT; width at half the PH), relative CSF volume (rCFV; area under the curve), and relative CSF flow (rCFF; rCFV divided by MTT). We determined how these metrics varied by age.

### Statistical analysis

Differences in CSF outflow metrics in age groups was determined using t-tests. Prior to the analysis, the data was tested for normal distribution using Shapiro-Wilk test and through visual inspection of Q-Q plots. Pearson correlation analysis was performed to analyze the correlation between relative CSF flow and age. All statistical analysis was performed using IBM SPSS v26.

### Validation study to distinguish signals of CSF from blood

We performed the following to ensure that our tagged MRI signal was from the CSF, and not the blood. First, we applied an oblique tag pulse perpendicular to the coronal imaging plane to avoid tagging the blood signal from the superior sagittal sinus (SSS) (as shown in supplemental Fig. S1). The SSS venous blood T2 value has been reported to be about 62 ms at 3T (*Lu et al., 2008*), depending on oxygenation and hematocrit levels (*Zhao et al., 2007; Chen et al., 2009*). To confirm if any contribution of blood signal occurred in our CSF measurements, we also acquired images with a long effective TE (TE_eff_) of 300 ms, at which no venous blood would show signal, and compared with images acquired at TE_eff_ of 30 ms. With TE_eff_ of 300 ms, only the long T2 component of CSF would be observable.

## RESULTS

To acquire detailed brain anatomy and to identify PSD and AG at the meninges, we performed high resolution T2-weighted 3D cSSFSE and T2 FLAIR acquisitions (Fig. 2). CSF in the SAS appears hypointense on FLAIR and hyperintense on 3D cSSFSE, whereas PSD demonstrates intermediate hyperintensity on both the FLAIR and 3D cSSFSE images (Fig. 2A). Note that multiple hyperintense signals, which are hypointense on FLAIR,(Fig. 2A), are seen in the 3D cSSFSE images, localized within the PSD and lateral SSS walls (Fig. 2B). These images were then used as locator images to identify PSD and to mark an ROI in the 3D coronal non-contrast MRI CSF outflow acquisition with varying inversion time (TI) periods. For visualization purposes, 3D cSSFSE images were fused with CSF tagged MR signals.

**Fig. 2.**
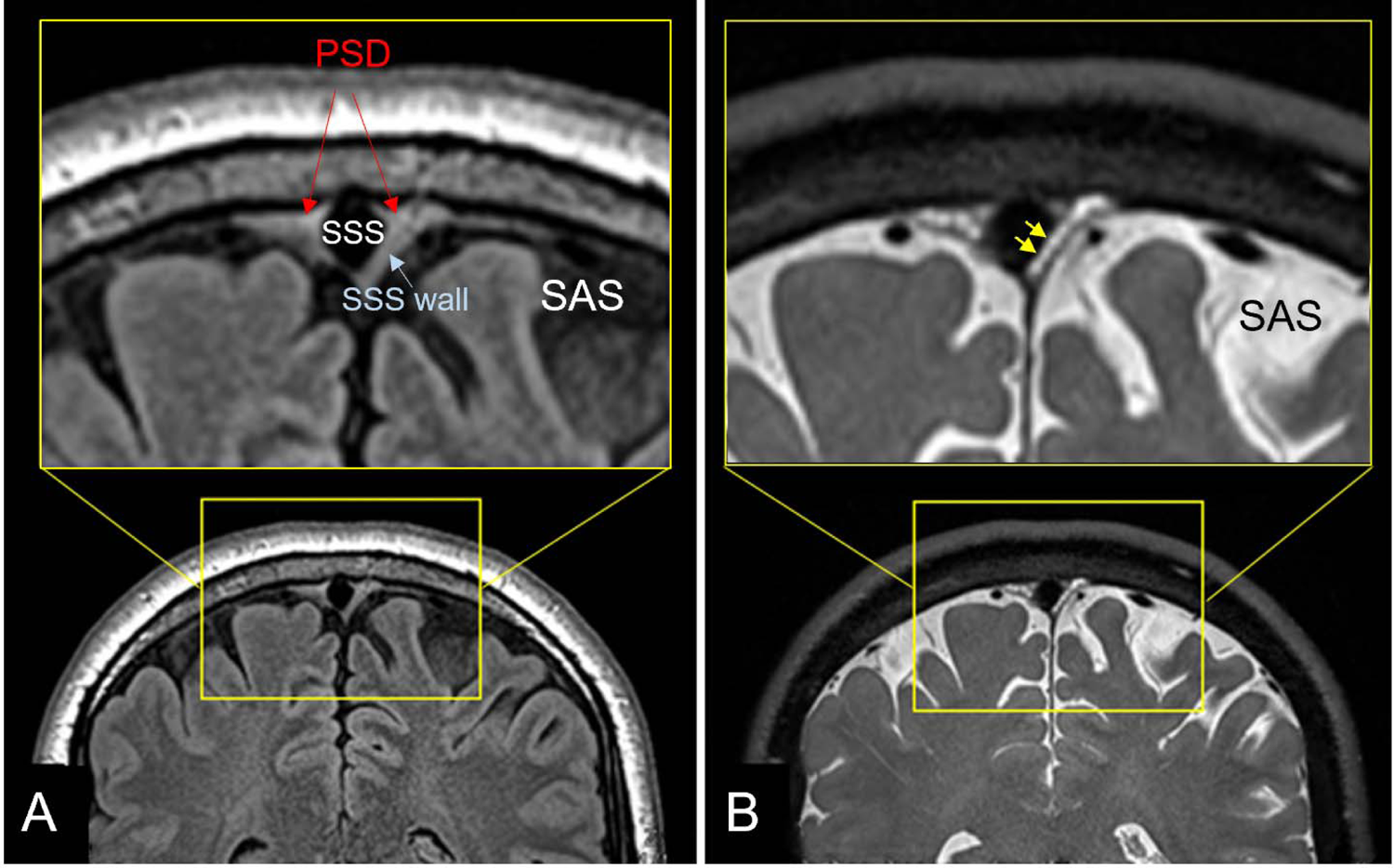
T2-weighted FLAIR and T2-weighted 3D cSSFSE of a healthy male adult, age 42. Top enlargement of the yellow box. **A)** FLAIR image depicts hypointense cerebrospinal fluid (CSF) in the subarachnoid space (SAS) and intermediate hyperintense parasagittal dura (PSD). **B)** 3D cSSFSE image depicts hyperintense CSF in SAS and intermediate hyperintense PSD and the lateral walls of superior sagittal sinus (SSS). Bright circles within the lateral wall may be arachnoid granulations (AGs) with bright signals, as shown in yellow arrows. Both images show hypointense blood vessels.

For CSF outflow imaging, we acquired coronal fluid spin labeling MR images with tag-on and tag-off acquisition and subtraction at varying TIs, as shown in Fig. 3. By acquiring both tag-on and tag-off images, the subtraction of tag-off from tag-on images shows movement of the tagged fluid out from the tag pulse area. Increasing the inversion recovery time (TI) period allows the tagged fluid to travel greater distance; however, the signal eventually decreases with longer TIs. Detailed explanations of sequence and spin states of tag-on and tag-off acquisitions are described in the Materials and Methods. To elucidate the egress pathways of CSF, we applied a simple subtraction method of time-resolved images by varying the TI periods, in which distance traveled by the tagged fluid varies by the TI periods. Fig. 4A shows the tag-off, tag-on, and the subtraction images with enlargement. The subtraction image presents the tagged CSF in the dura mater, PSD and alongside the SSS lateral wall. Fig. 4B shows the subtracted color images of the same slice using TI of 500, 750, 1000, 1250, 1500, and 2000 ms of the same center slice at Z = 0 mm, as time-resolved images. This allows visualization of the tagged CSF as it moves out from the tag at the dura mater toward the PSD at TI of 750 ms, and straight down to the lateral wall of SSS located closer to the tag pulse at TI between 1000 and 1500 ms. The signal decreases at TI of 2000 ms. Fig. 4C presents the color images of contiguous slices covering about −1.5 to +1.5 mm (3 mm), showing the tagged CSF signals from dura mater to the lateral wall of SSS in all 3D slices. We also observed an additional signal extending directly from the SAS toward the SSS, as indicated by the red arrow. In our subtracted images, we observed the CSF from dura mater or PSD to the lateral wall in most participants, although the signal was weak in some. Fewer participants exhibited signals from the SAS to the SSS. Supplementary Figure 2 shows example subtracted color images from 3 participants, demonstrating tagged signals appearing to move out from the dura mater from all participants, albeit with varying degreess of signal strength.

**Fig. 3.**
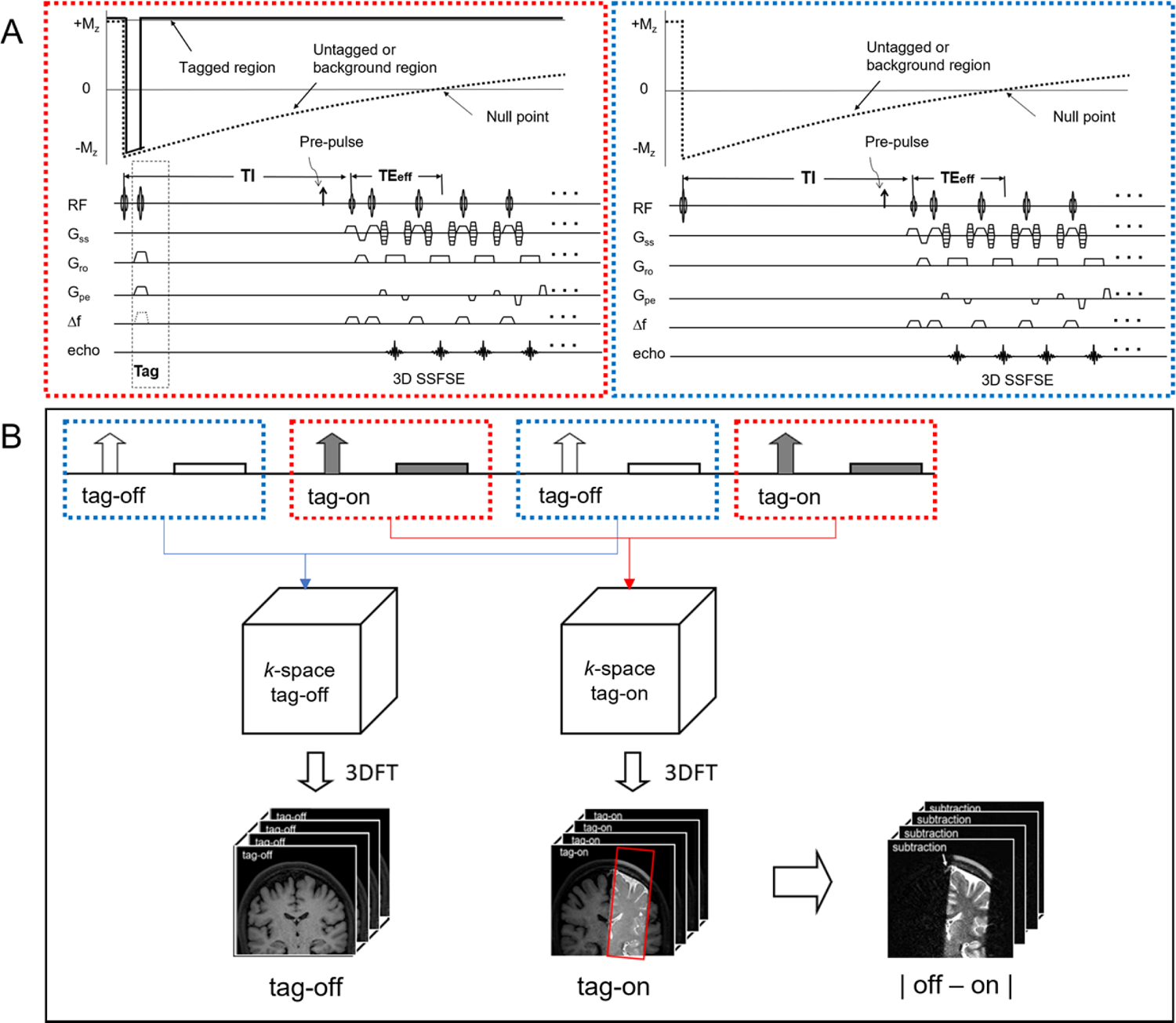
A) Pulse sequence diagram of tag-on and tag-off 3D acquisition using 3D cSSFSE with magnetization states. Each slice encoding of tag-on and tag-off acquisitions is alternately acquired. Pre-pulse such as SPAIR fat suppression pulse can be applied before the readout acquisition. **B)** Post-processing of 3D Fourier-transformed data of tag-off and tag-on brain images with a rectangular tag position in red (images at TI of 1500 ms, image details in **Fig. 4**). Simple subtraction enables visualization of movement of the tagged signal out of the tag region.

**Fig. 4.**
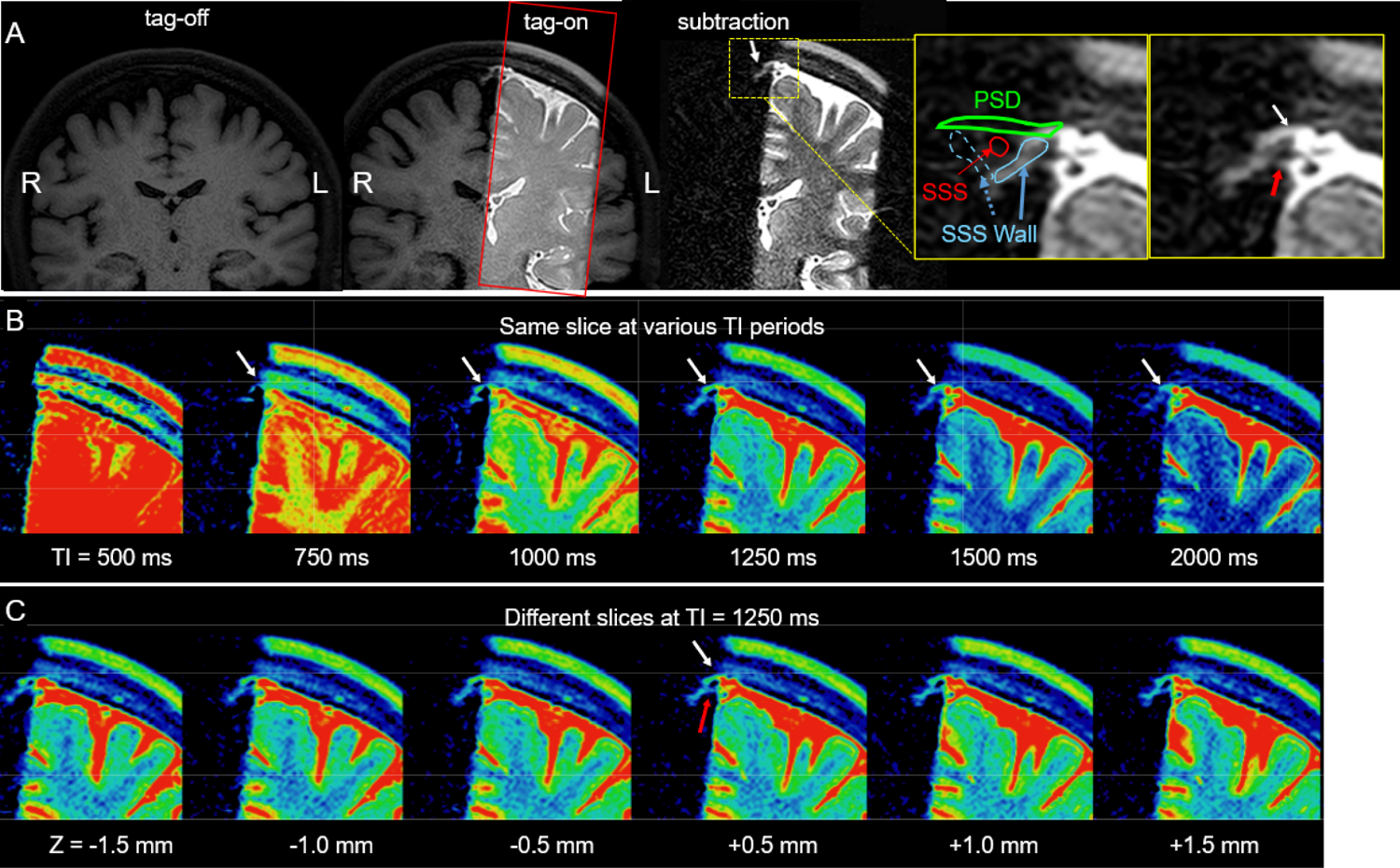
A) Images of tag-off, tag-on, subtraction, and the enlargement of the subtraction obtained at TI of 1500 ms. The Time-SLIP pulse is shown in the red rectangular box. Enlarged images with and without labels show fluid moving out from the tagged region (left brain hemisphere) to the parasagittal dura (PSD) (white arrow) extended from the dura mater and the lateral wall of the superior sagittal sinus (SSS) (red arrow). Segmented labels show parasagittal dura (PSD in green), lateral walls of superior sagittal sinus (SSS) (light blue) (left wall in a solid line and the right wall in a dotted line), and SSS in red. Note that the signal by the red arrow shows a possibility of a second pathway of CSF flow from the subarachnoid space (SAS) to the SSS, via the left lateral wall of SSS. Note that there are no noticeable signals at the right PSD and lateral wall of SSS from the right brain hemisphere consistent with no Time-SLIP pulse applied to this hemisphere. **B)** Color map of subtracted images from the center slice at Z=0 mm location of 3D images at TI of 500, 750, 1000, 1250, 1500, and 2000 ms. The tagged CSF moved out to PSD at 750 ms, straight down to lateral wall of SSS at TI of 1000 to 1250 ms, and gradually decreased in signal intensity by TI of 2000 ms. **C)** Contiguous slices of subtracted color maps of 3D volume images obtained at TI of 1250 ms. Note that contiguous slices show marked movement of CSF toward the lateral wall of SSS in every slice (or a 3 mm Z direction distance) of 3D volume images.

Next, we quantified the CSF flow using signal increase ratio (SIR) of the tagged CSF MRI in the SSS region. The calculation and post-processing of SIR are explained in the Materials and Methods. Fig. 5A shows the coronal image and tag pulse (red). Fig. 5B shows enlarged oblique fusion images of 3D SSFSE and tagged MRI signals at the ROI covering the SSS, the wall of SSS, and PSD in various TI periods. Bright white signal indicates strong tagged fluid signals, with an intensity increase up to 300%. At increasing TIs, this fusion technique captures snapshots of the egress pathway of CSF from dura mater into SSS, alongside the wall of SSS, visualized in oblique 3D images (Fig 5B; video movie M1). Magnified straight coronal views in Fig. 5C demonstrate tagged CSF at the level of the PSD entering the lumen of the SSS from 500 to 1000 ms. Fig. 5D shows quantitative measures of CSF signals at each TI period. When applying one tag pulse to the left hemisphere, the tagged CSF signal travels to the SSS, reaching a peak flow, and disappearing with longer TI periods. By systematically increasing the TI period, and using tag-on, tag-off subtraction, we can visualize and quantify the path travelled by tagged-CSF until the signal disappears with long TI periods. After the non-selective IR pulse, which is applied to both the tag-on and tag tag-off acquisitions, the background signals outside the tag region experience the same exponential T1 relaxation magnetization and are cancelled out by the subtraction; only the tagged signals remain visible. Thus, venous blood signals in the SSS are likely to be cancelled out by the subtraction of tag-on and tag-off at each TI. To further avoid SSS venous contamination, the spin labeling tag-on and tag-off images were acquired in the coronal orientation while the tag labeling pulse for endogenous CSF tracing was applied in the oblique sagittal orientation parallel to the long axis of the SSS.

**Fig. 5.**
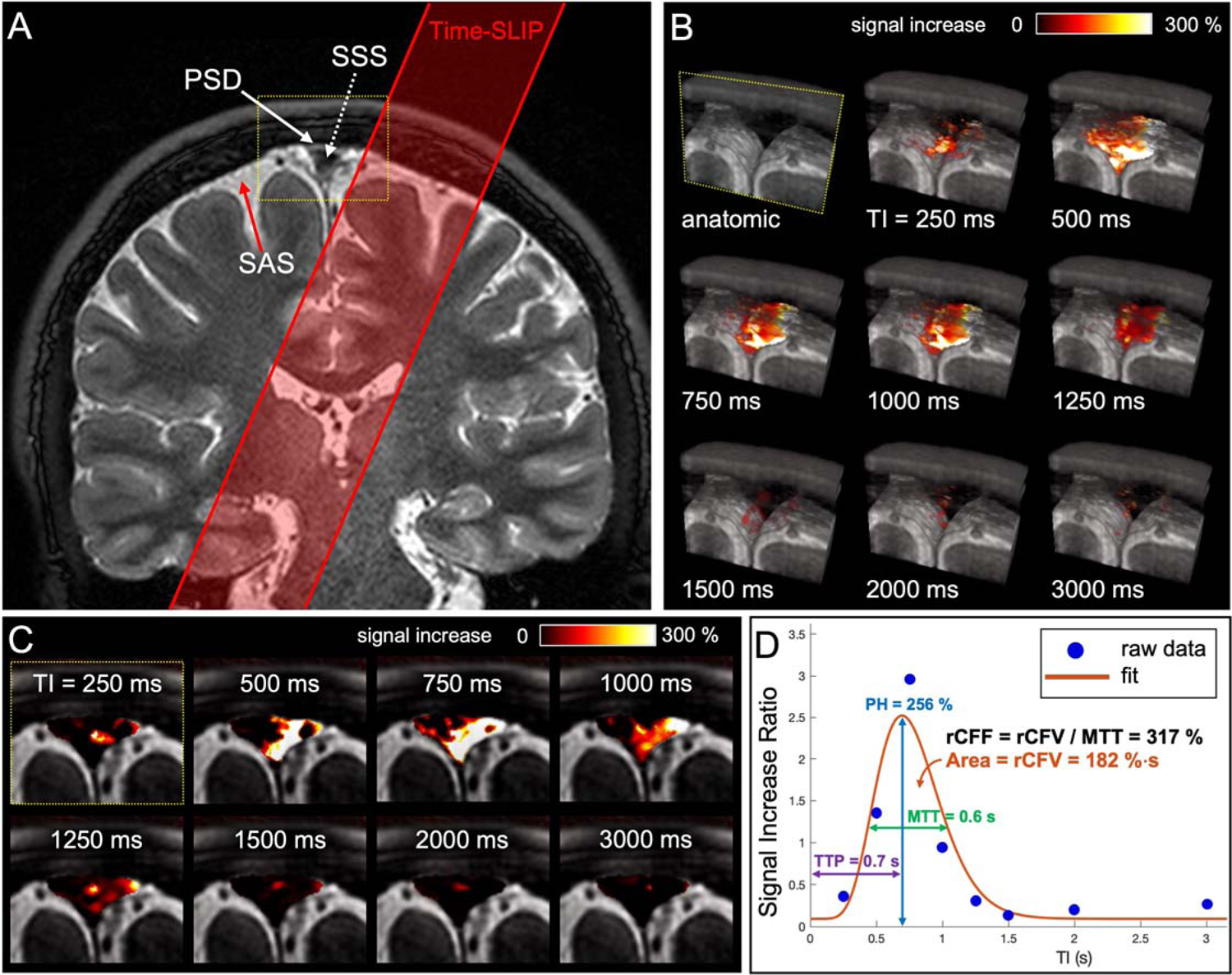
A) Coronal 3D cSSFSE image with a spin labeling tag pulse (red box). The Time-SLIP tag pulse is placed about 10 mm away from the superior sagittal sinus (SSS). **B)** A series of oblique 3D spin labeling images with various inversion times (TIs) fused over 3D cSSFSE image, enlargement of yellow box in A). Note that the tagged signals increase from the Time-SLIP on the left brain at the parasagittal dura (PSD) region at TI of 500 ms and then disappear or egress into the SSS by 1500 ms. **C)** Straight coronal fusion images of tagged MRI signals around the SSS region over 3D cSSFSE images at various TIs show tagged fluid outflow from the tag pulse to dura mater and PSD into the SSS. In addition, the left lateral wall is highlighted around TI of 500 ms and 700 ms **D)** Tagged CSF outflow signal at the SSS. Circles show the data points and the line indicates the curve fit. Peak height (PH) is 256%, mean transition time (MTT) is 600 ms, time to peak (TTP) is 700 ms, relative CSF volume (rCFV) is 182%·sec under the curve, and relative CSF flow (rCFF) is 317%, obtained by dividing rCFV by MTT.

To quantify CSF outflow metrics, we curve-fitted CSF MRI outflow signal vs. TI data (Fig 5D). The peak of the signal is seen at TI around 700 ms, reaching a peak height (PH) of 256%. The signal returns to the baseline after TI of 1500 ms. The relative CSF flow (rCFF) is 317%.

To determine the effect of age on the outflow of tagged CSF measures, we imaged 16 healthy participants ranging in age from 19 to 71 years. Supplementary Figure 3 shows typical time-resolved images of two younger and two older adults with enlarged fusion images of 3D SSFSE and tagged MRI signals at the ROI covering the PSD, SSS walls, and SSS, in various TI periods. The younger adults display greater intensity signals than the older adults. Fig. 6A shows the quantitative CSF outflow of the entire SSS region by participant age. Although our sample size is small, there was a strong negative correlation of rCFF with age (*r* = −0.82; *p*<0.001). This relationship appears to be non-linear, with greater decline of rCFF among those over 60 years. Thus, we categorized participants into younger (19 to 59 years, n=8) and older (60 years and older, n=8) groups to better assess age differences. Fig 6B shows group differences in tagged CSF outflow curves. Fig 6C shows group differences in quantitative outflow measures. PH, rCFV, and rCFF were significantly lower among older than younger adults (each *p* < 0.01). Time to peak (TTP) and mean transit time MTT did not differ between groups.

**Fig. 6.**
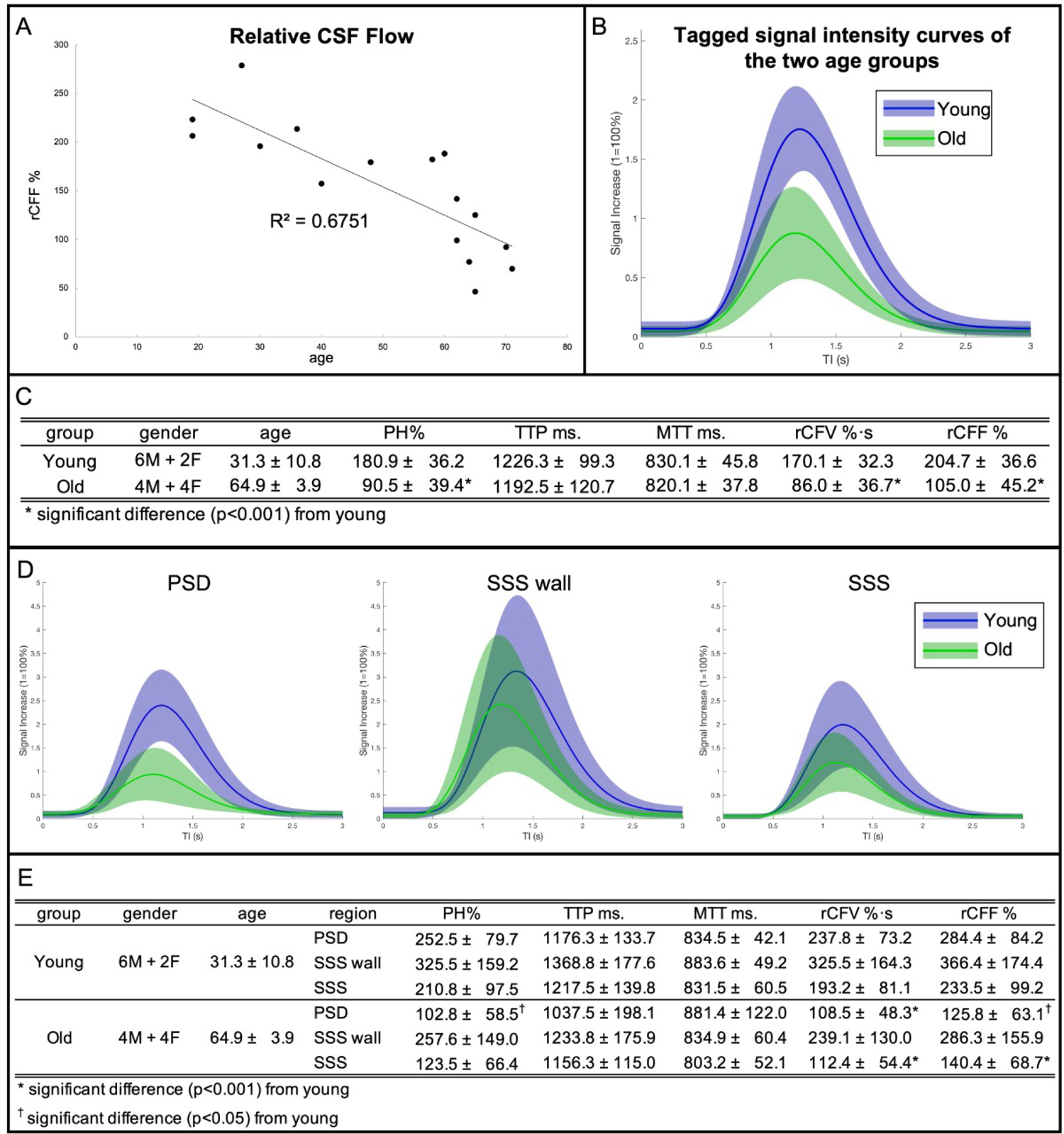
A) CSF outflow of the entire SSS region on 16 participants between 19 and 71 years show a decline of outflow with age, especially in those over 60 years of age. **B)** Tagged fluid outflow rate of two groups: younger (19-59 years) and older (60-71 years) adults. The amount of outflow signal is markedly decreased in the older group, as compared to the younger group. **C)** Quantitative CSF outflow measures between the younger and older groups. Tagged CSF outflow measures are peak height (PH), relative CSF volume (rCFV), and relative CSF flow (rCFF) for each age group and for all participants. The time-to-peak (TTP) and mean transit time (MTT) values were similar in the two groups; however, PH, rCFV, and rCFF were significantly different between the two age groups. **D)** Individual ROI analysis of PSD, SSS wall, and SSS in younger and older age groups and their quantitative measures (**E**). PSD shows a significant difference in PH, rCFV, and rCFF between younger and older groups. SSS presents a significant difference in rCFV and rCFF between younger and older groups. Note that the TTP is similar between PSD and SSS, but slightly longer in SSS whereas TTP of the lateral SSS wall is longer than that of PSD and SSS.

Next, for all 16 participants, we segmented the SSS region into the 3 ROIs of PSD, SSS wall, and SSS shown in the top right panel of Fig. 4. Fig. 6D shows the tagged CSF outflow curves of these three regions in the younger and older groups. Fig. 6E shows the corresponding tagged CSF metrics. Quantitative outflow measures of PH, relative CSF fluid volume (rCFV), and rCFF in the PSD and SSS were significantly lower among the older group. However, there were no differences in these values for the SSS wall. Note that the difference in TTP between PSD and SSS, with slightly elevated TTP in SSS relative to PSD, is consistent with the direction of flow from the PSD into SSS (see Fig 1). However, TTP of the lateral SSS wall is longer than that of PSD and SSS, suggesting a different pathway from that implied by the time course of PSD to SSS.

For reproducibility, we acquired two quantitative measures of tagged CSF measures consecutively in one scan session for two participants (aged 48 and 62 years). Results are shown in Supplementary Figure 4. PH was the most reliable measure, with percent differences between the two scans of < 2% and <7% for the two participants. For other flow parameters rCFF and rCFV percent difference was not more than 23%.

In addition, we examined whether there was any signal contribution from blood in our tagged MRI measurements. The source images of tag-on and tag-off images in various TI images on all 16 participants obtained using TE_eff_ of 30 ms were reviewed by a medical doctor (XZ). We did not detect any bright blood signals in the SSS and the surrounding blood vessels such as cortical veins, as shown in Fig. 4, Fig. 7A-D, and Supplementary Figure 3. Based on our sequence design, bright signal within the Time-SLIP pulse moves out as bright while the dark signal moves out dark. At each TI period, the background signals experience an exponential return in both tag-on and tag-off acquisitions, as shown in Fig. 3. The tagged region also showed dark signal intensity in the regions corresponding to blood vessels; therefore, it could be deduced that no blood signals from the tagged pulse contributed to our tagged MRI measurements. Only bright signals (+Mz) in the tag region contribute to the tagged MRI signals, as indicated in Fig. 3A.

**Fig. 7.**
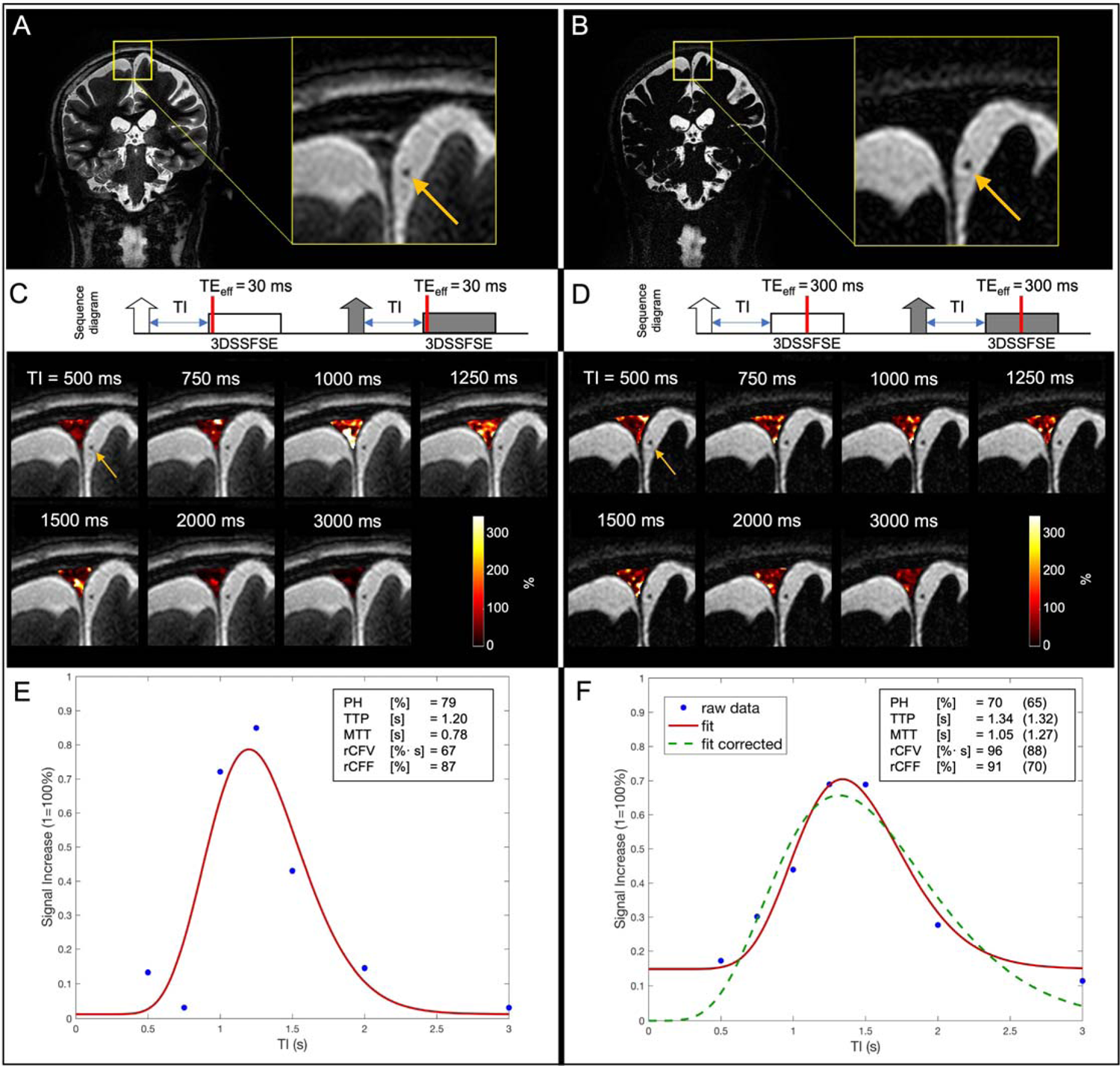
The tag-on and tag-off acquisitions using an effective TE (TE_eff_) of (A,C,E) 30 ms and (B,D,F) 300 ms. A and B are coronal images and their enlarged anatomic images of TE_eff_ of 30 ms and 300 ms, respectively. C and D show the acquisition sequence showing the contrast-determined time of TE_eff_ of 30 ms and 300 ms by red line, and their various inversion time (TI) images, respectively. The fusion image with TI 500 ms was used with tagged MRI color images covering the entire SSS region including PSD, SSS walls, and SSS at a slice of 3D images using various TI periods. Both TE_eff_ of 30 ms and 300 ms show tagged CSF signals at the PSD, SSS walls, and SSS, while showing dark blood signals (yellow arrows) throughout TI periods. Tagged MRI signal curves of both TE_eff_ of (E) 30 ms and (F) 300 ms show similar tagged MRI curves; tagged MRI signal increase (wash-in) and decrease (wash-out) were both around 1100 to 1300 ms, with similar PH. However, mean transit time (MTT) was a bit longer, relative CSF flow (rCFF) and relative CSF volume (rCFV) were higher in data obtained at TE_eff_ of 300 ms, as compared to TE_eff_ of 30 ms. MTT was increased by approximately 270 ms, which is the difference between TE_eff_ of 300 ms and 30 ms. This may be due to adding an extra contrast-determined time of TE_eff_ to the TI period. During the TE_eff_ of 300 ms period, tagged signals within the 3D slab may travel in multiple directions to increase tagged MRI outflow measures. Regarding the baseline of (F) TE_eff_ of 300 ms, it is a non-zero baseline due to the lower signal but higher contrast (more T2-weighted) at TE_eff_ of 300 ms, as compared to TE_eff_ of 30 ms. From the Equation [1], lower signal in the region of interest due to the long TE_eff_ likely to introduce an extra signal bias like a ratio of small signals in both numerator and denominator. To correct this bias, we have added an extra data point (SIR = 0% at TI = 0 ms) based on the assumption that no signal is present at TI = 0 ms. Dotted green line shows the corrected curve fit with zero starting point with the corresponding tagged fluid metrics in the parentheses.

Lastly, to ensure that only CSF and not blood is imaged, we scanned increasing TE_eff_ in 3D SSFSE readout to 300 ms, to selectively acquire long T2 components of CSF, rather than observing surrounding blood signals (shorter T2) that would have decayed. The SSS venous blood has T2 value of 62 ms (*Lu et al., 2008*); therefore, there should not be blood contribution at TE_eff_ of 300 ms. Fig. 7A and B show the coronal images with enlargements obtained using TE_eff_ of 30 and 300 ms. Fig. 7C and D show the simple sequence charts of tag-on and tag-off acquisitions using TE_eff_ of 30 ms and 300 ms, as well as fusion images at each TI period. The contrast-determined time is the TE_eff_ period in 3D SSFSE, as indicated by red bars. The fusion images of Fig. 7C and D show the background tag-off images using TE_eff_ of 30 ms and 300 ms at TI of 500 ms with tagged MRI signals covering PSD, lateral walls of SSS, and the SSS. Note that the background images show that the PSD region is bright in TE_eff_ of 30 ms, but dark in TE_eff_ of 300 ms, due to the T2 difference. The color maps of the entire PSD region show the tagged MRI signals from the lateral wall of SSS with bright white color, indicating high values. Fig. 7E and F show the results of tagged CSF curves obtained using TE_eff_ of 30 ms and 300 ms. Both TE_eff_ of 30 and 300 ms produce similar quantitative metrics of PH, rCFV, and rCFF. Note that TE_eff_ of 30 ms and 300 ms provide a similar TTP with a reasonable delay due to an extra time of TE_eff_ of 300 ms, consistent with an increased TE_eff_. However, the MTT was slightly longer and the baseline was higher in the TE_eff_ of 300 ms, as compared to TE_eff_ of 30 ms. We applied the |tag-off – tag-on|/|tag-off| images as an SIR in our tagged CSF signal analysis because the TE_eff_ of 300 ms results in a low signal-to-noise ratio (SNR) as compared to TE_eff_ of 30 ms in both tag-on and tag-off images, resulting in a low SNR. This low SNR may be amplified by dividing by the low SNR in the tag off image. We also added the calculation using a TI of 0 ms, for which zero signal was assumed. This experiment confirmed that both TE_eff_ of 30 ms and 300 ms detect CSF outflow in the SSS region and not blood.

## DISCUSSION

The findings from the time-resolved spin-labeling MRI study demonstrate, for the first time, a method to directly image the intrinsically tagged CSF outflow without administration of exogenous tracers or contrast agents in healthy adults. By utilizing spin-labeled MRI as an endogenous tracer at the tag region of dura mater and SAS, we were able to visualize tagged CSF in two pathways: from dura mater to PSD and SAS into the SSS and directly from SAS to SSS. The small differences in TTPs between PSD and SSS are consistent with flow of the tagged CSF from dura to PSD and then to SSS, while the slower TTP of SSS lateral wall is consistent with a direct pathway from SAS to SSS, likely via AGs (*Pollay 2010; Sakka et al., 2011*). This pathway was best visualized in the subtracted color images from young adult participants, many of which showed signal extending into the SSS lateral wall from the SAS about a few millimeters below the PSD pathway. Although our techniques utilize ungated 3D cSSFSE with slice encoding per every TR of 5400 ms, we nevertheless observed that all slices in the 3D volume demonstrated the outflow of CSF. This implies that outflow may occur constantly in the brain as a steady state, as has been previously suggested (*Metzger et al., 2017*).

Outflow of CSF containing waste products including Aβ is essential for the maintenance of brain health (*Benveniste et al., 2019; Hou et al., 2019*). Aging is one of the major risk factors of neurodegenerative diseases (*Hou et al., 2019*). Murine models have shown reduction in glymphatic clearance with age (*Ma et al., 2017)*. Our non-contrast MRI study also demonstrated a decline of CSF outflow with age in healthy adults. Our method, which is suitable for use in anyone without MRI contraindications, may facilitate establishment of normative values for CSF outflow. This may allow further investigation of the role of CSF outflow in neurodegenerative disorders and potential advancement of a novel biomarker predictive of risk for the development of dementia.

The present technique exploits the intrinsic image contrast between the SAS and compartmentalized components of CSF at the PSD, demonstrating low and high signal intensities on FLAIR and 3D cSSFSE, respectively. This finding is supported by recent work demonstrating the presence of several distinct components of CSF signal in the brain based on T2 or spin-spin relaxation times (2000 ms at the CSF of SAS and T2 of 400 ms for the PSD) using non-negative least squares in pixel-by-pixel analysis (*Oshio et al., 2021)* and most recently by comparison of signals reflecting various albumin concentrations with results from human imaging (*Albayram et al., 2022*). Additional work further helped compartmentalize these signals using FLAIR imaging^7^ to demonstrate the complex, sub-millimeter cross-sectional structure of the PSD extending along the lateral wall of the SSS (*Aspelund et al., 2015; Louveau et al., 2015; Absinta et al., 2017*). Given the close opposition of the PSD and the SAS, and their intrinsic differences in spin-spin relaxation times, an active CSF-mediated exchange of molecules between the brain tissue and dural lymphatic vessels has been postulated as the structural underpinnings of the glymphatic system (*Ringstad et al., 2020*).

Results of our validation study suggest that tagged MRI signal acquired using the present technique likely stems from the longer T2 components of CSF, not from the blood. We have shown that there is no bright blood signal in either the tag-on or tag-off source images, in the SSS and surrounding vessels. The tagged fluid flow-out technique shows only the tagged region with bright signals move out bright, while dark signals move out dark (*Yamada et al 2008; 2013*). We also compared images acquired at both short (30 ms) and long (300 ms) TE_eff_. Because of the short T2 of approximately 62 ms of SSS venous blood, the blood signal would experience marked decay in the long TE_eff_ images (*Lu et al., 2008*). The TE_eff_ of 30 and 300 ms resulted in similar tagged MRI signal curves with similar PH, rCFV, and rCFF, which would not be expected if blood contributed to the signal. The longer MTT in TE_eff_ of 300 ms can be explained by the tagged fluid within the 3D excitation slab during the TE_eff_, which allows fluid to move in multiple directions in PSD, causing an elongation of tagged signals returning to the baseline, as shown in Fig. 7. Taken together, these findings support that our non-contrast spin-labeling MRI technique enables assessment of the flow of CSF rather than blood.

Regarding the tagged CSF traveling time, interestingly, the mean outflow time of endogenous CSF from the dura mater to SSS using this technique averaged less than a couple seconds, rather than on the order of hours or days as previously reported with the use of intrathecal administration of GBCA (*Ringstad et al., 2020*). This may reflect the relatively limited diffusivity of GBCA across the selectively impermeable barrier of the PSD. In addition, another study with intravenous administration of GBCA showed immediate enhancement at cortical veins (*Naganawa et al., 2020*). This suggests subpial fluid flow around the cortical veins may be responsible for fast turn over. However, unlike tracer studies, we observed the egress pathways of intrinsic CSF outflow from PSD to the SSS and SAS to SSS within a couple of seconds. Therefore, our study may fill the technical gap between the GBCA-tracer MRI and intrinsic CSF outflow.

We obtained preliminary data on the reproducibility of our method on 2 participants. There was excellent reliability of PH and TTP in both participants; however, one of the participants showed large percentage differences in rCFV and rCFF, mainly due to high variability in MTT. The MTT metric strongly depends on the temporal resolution. Thus, reliability can likely be improved by increasing the data sampling to improve the temporal resolution which will be assessed this in future studies.

Our time-resolved CSF MRI has the following limitations: temporal and spatial resolution of 4D acquisition is about 250 ms around the peak flow with submillimeter resolution; the present study also applied tagging only one side of brain hemisphere at the top of the brain; small sample size; and the SIR baseline of tagged MRI signal curve relies on the TE_eff_ of the tag-on images as non-zero signal. Nevertheless, our time-resolved study shows oblique-sagittal tagged CSF travels both pathways from the dura mater and PSD to the SSS and SAS to SSS, via the lateral wall of SSS by the subtraction color images and the quantitative SIR analysis. Regarding the extra selective IR pulse in the tag-on acquisition, there is a magnetization transfer (MT) effect in the brain parenchyma; however, we believe CSF has less MT effect.

## CONCLUSION

In conclusion, our experimental findings from the subtraction of tag-on and tag-off images in the healthy human adults reveal egress pathways of tagged CSF outflow from PSD to the SSS and from SAS to SSS. This outflow is quantifiable, and initial results show strong differences in CSF outflow metrics with age among healthy adults. Our non-invasive non-contrast tagged MRI method may be an important enabling technology to non-invasively study CSF outflow in sleep, across the circadian rhythm and in response to exercise and other interventions that may impact the glymphatic system. Furthermore, this method may help increase knowledge of CSF outflow contributions to pathological brain changes in aging and neurodegenerative disorders.

## Abbreviations

AG: arachnoid granulations

CSF: cerebrospinal fluid

cSSFSE: centric *k*_y_–*k*_z_ single-shot fast spin echo

FLAIR: fluid-attenuated inversion recovery

FOV: field of view

GBCA: gadolinium-based contrast agent

MT: magnetization transfer

MTR: magnetization transfer ratio

MTT: mean transit time

PSD: parasagittal dura

PH: peak height

ROI: region of interest

rCFV: relative CSF fluid volume

rCFF: relative CSF fluid flow

SAS: subarachnoid space

SIR: signal increase ratio

SNR: signal-to-noise ratio

SSS: superior sagittal sinus

TTP: time-to-peak

## Funding

This work was supported by NIH grants RF1AG076692 (M.M.) and R01HL154092 (M.M.), and a grant by Canon Medical Systems, Japan (35938).

*The funders had no role in study design, data collection, and interpretation, or the decision to submit the work for publication.

The authors thank Professor Alexander Norbash, Department Chair of Radiology, for his support on this project. The authors also thank Dr. Xiaowei Zhang for reviewing the regions of interest and literature search, Ms. Sheronda Statum for recruitment of participants, Mr. Jirach Kungsamutr for his help on data analysis, Dr. Yoshimori Kassai and Mr. Yurian Falls of Canon Medical Systems for their technical support.

## Author contributions

**V.M.** Data collection, Acquisition methodology, Development of post-processing software, Data analysis, Writing-draft preparation**. W.B.** Data collection, Development of post-processing software, Data analysis, Visualization, Review and Editing. **X.Z**. Review of regions of interest for quantitative analysis, **A.Y.** Data collection, Initial investigation, Image review. **L.K.M:** Data interpretation, Writing – Substantial Revision. **M.A.M.** Image review, Review and Editing. **M.M.** Conceptualization, Development of pulse sequence design, Data collection, Initial investigation, Writing-Original draft preparation, Reviewing and Editing.

## Data and materials availability

All data associated with this study are present in the paper or in the supplementary material. None of the material has been published or is under consideration for publication elsewhere. We have deposited anonymized subject data (**DOI: 10.5281/zenodo.6792179**) and demo code at (**DOI: 10.5281/zenodo.6792214**).

## Supplementary Information

### Supplemental Figure and Captions

**Supplementary Figure S1.**
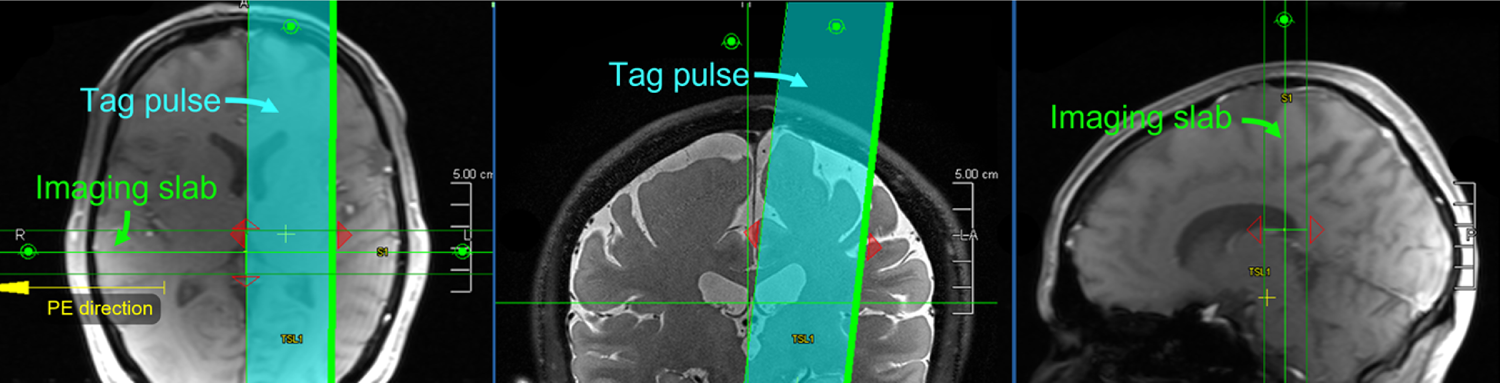
Axial (left), coronal (middle), and sagittal (right) localizer images from the left show the positions of the 3D imaging slab and the tag pulse. The oblique tag pulse is perpendicularly applied to the 3D coronal imaging slab.

**Supplementary Figure S2:**
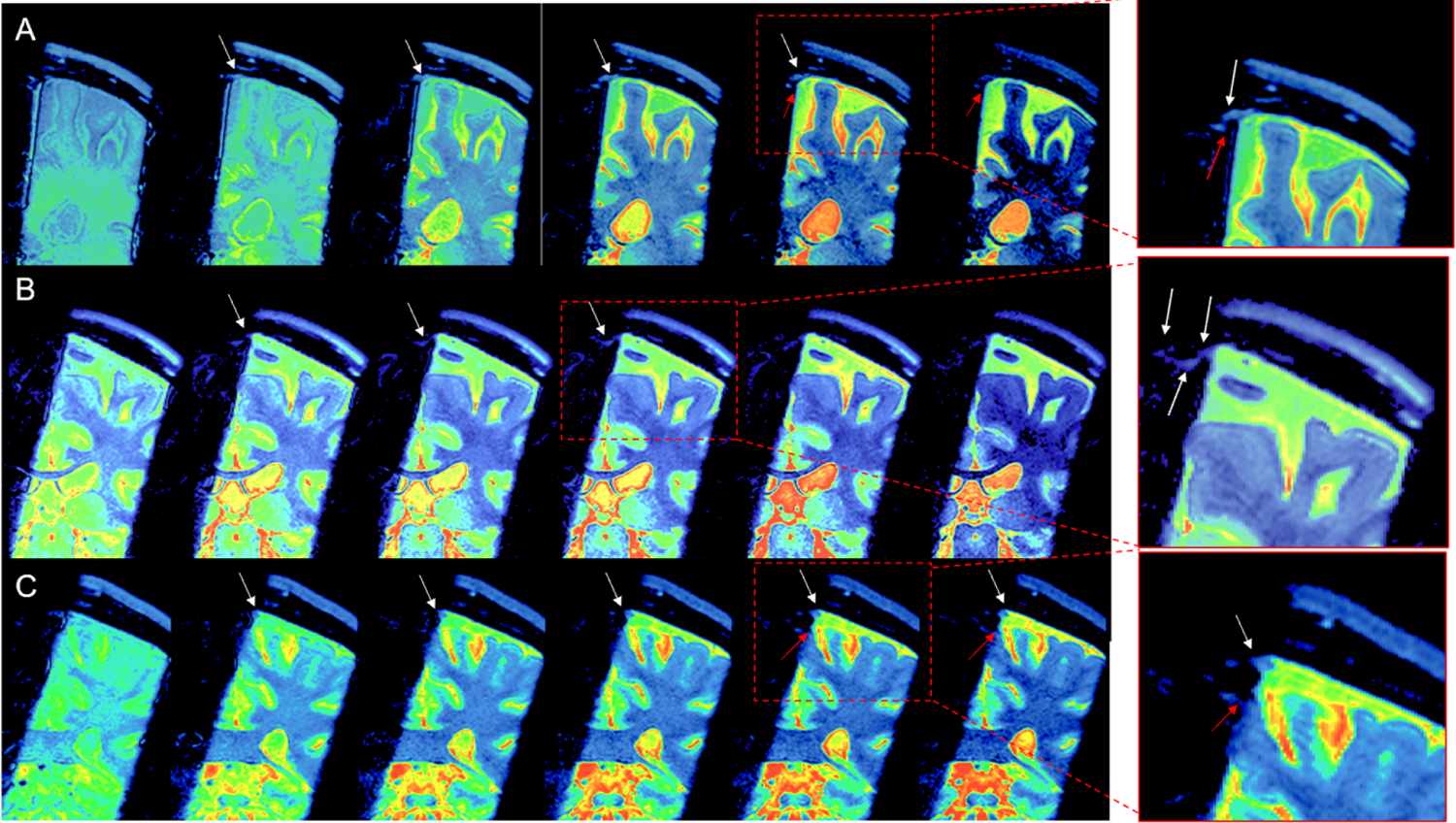
Three subtraction color images of tagged CSF at TI of 500, 750, 1000, 1250, 1500, and 2000 ms of A) 30 y.o. male, B) 19 y.o male, and 36 y.o. female participants. Enlargements show the CSF from dura mater (white arrows). A) and C) show a signal from another route from subarachnoid space (SAS) to the lateral wall of superior sagittal sinus (SSS) (red arrows). B) shows tagged CSF signals that seem to travel to the left lateral wall and then to the right lateral wall of SSS.

**Supplementary Figure S3:**
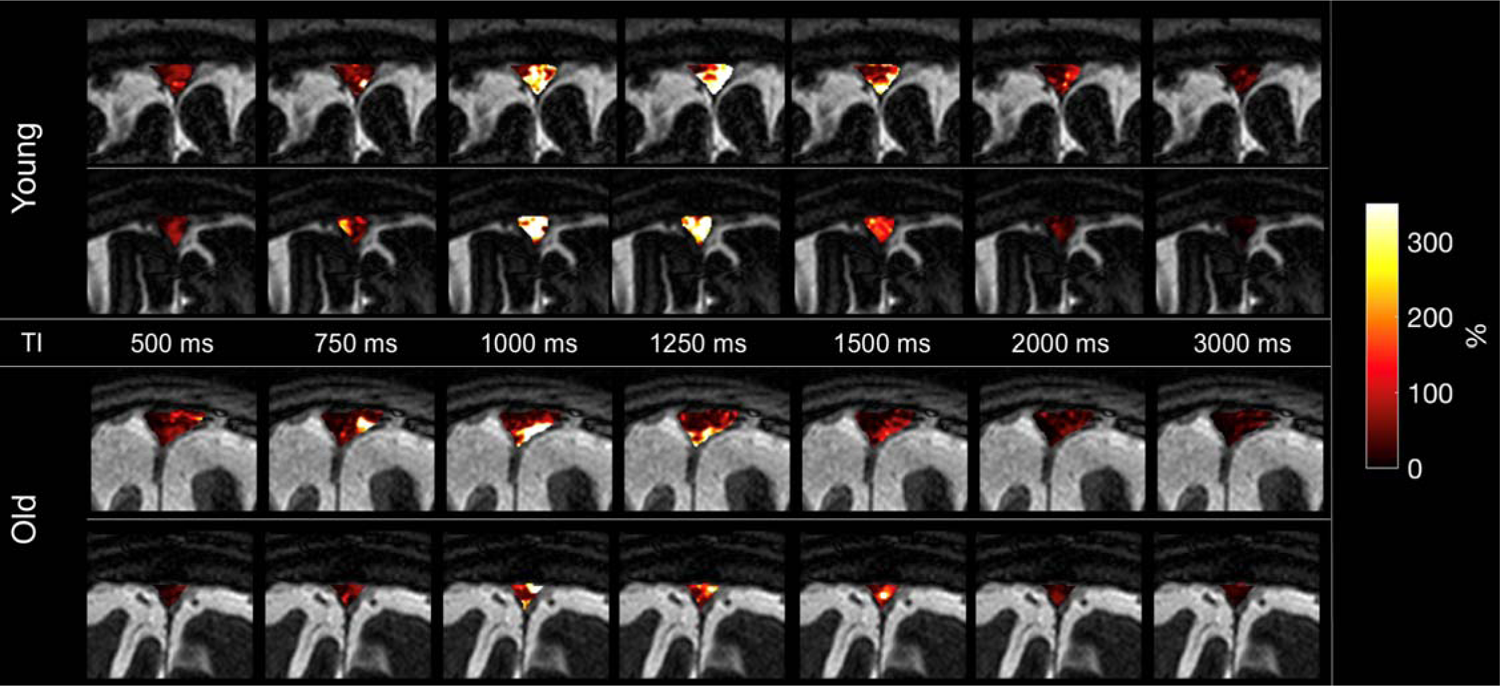
Four example images of tagged glymphatic color maps at TI of 500, 750, 1000, 1250, 1500, 2000, and 3000 ms in two younger (top) and two older (bottom) adults. The younger adults show bright signals in the SSS region 1000 to 1500 ms, signals in the older adult images are less bright.

**Supplementary Figure S4:**
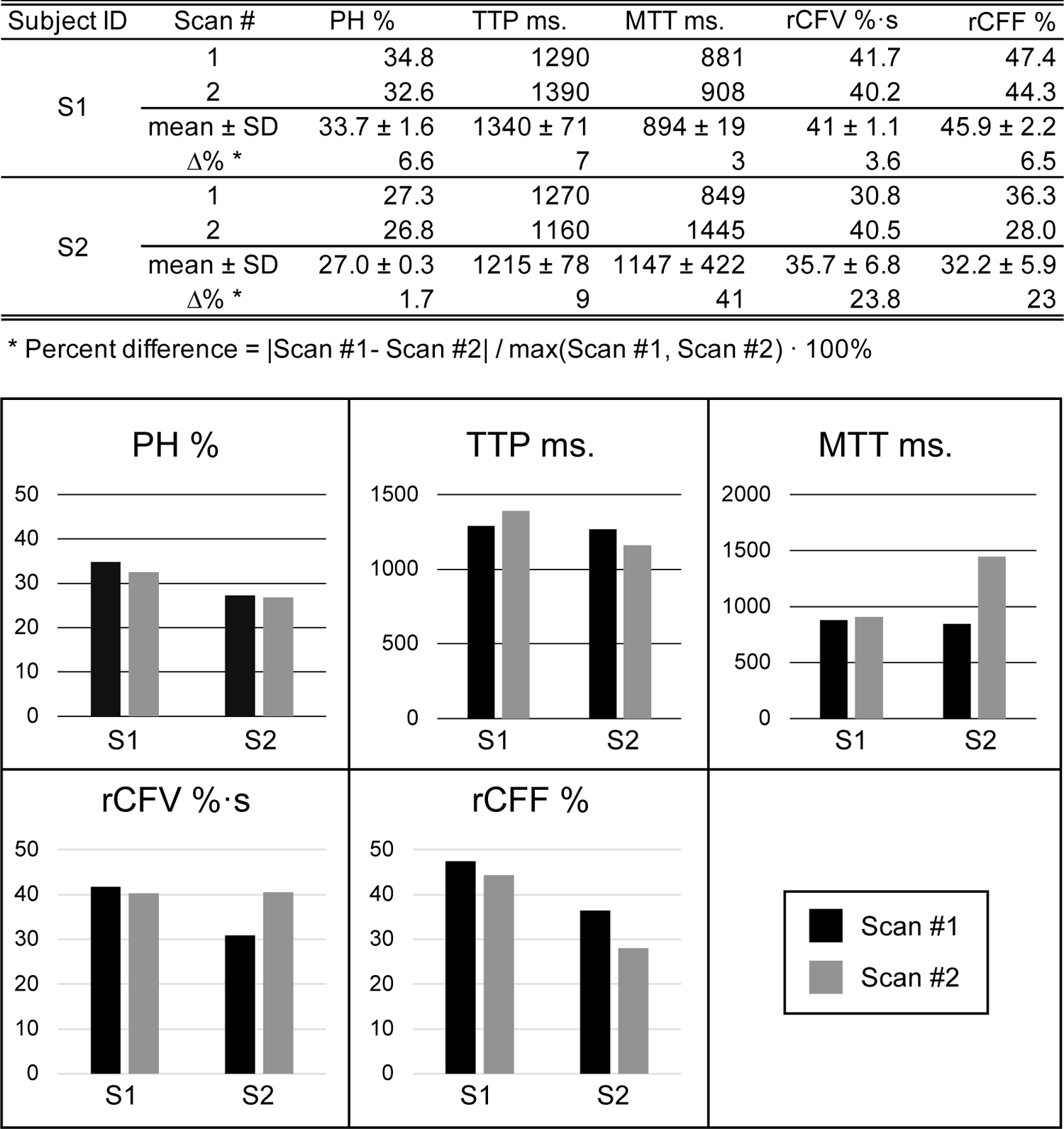
Reproducibility, Re-test reliability of the tagged CSF outflow measurements for two participants. Percent differences in peak height (PH) and time-to-peak (TTP) have less than 10%. For participant 1, there was little difference in outflow metrics between the two scans. For participant two, there was no difference in PH between scans, and little difference in TTP, but the larger difference in mean transit time (MTT) (41%) between the two scans resulted in somewhat more variability between scans for rCFV and rCFF (23%).

**Video of Fig. 5A and 5B.**
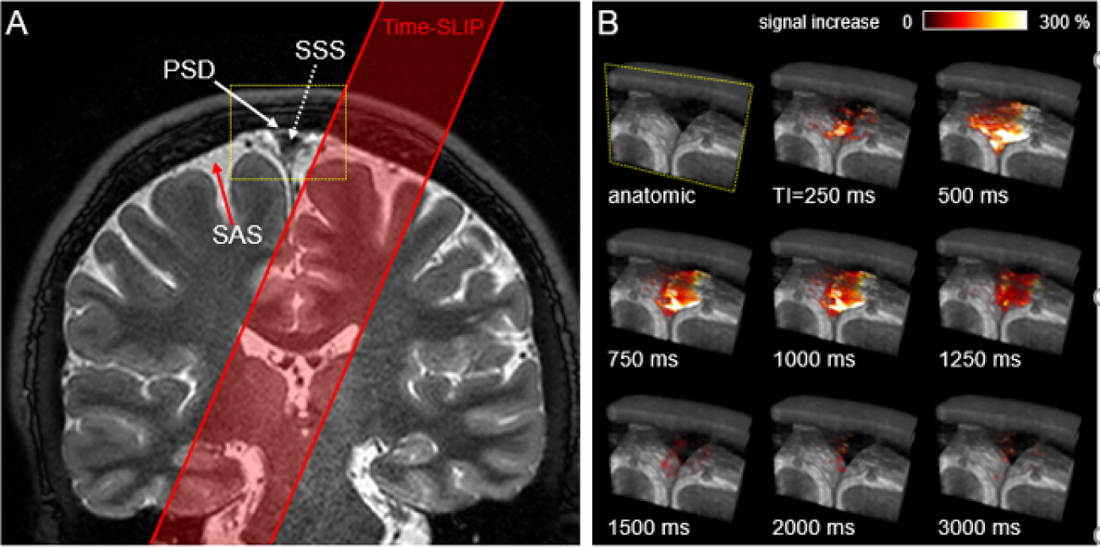
Left and middle images from Fig. 5 A and B, respectively. Coronal T2-weighted 3D cSSFSE image. The yellow box indicates the meninges region. Bottom video: Enlargement of yellow box. The oblique outflow 3D images covering 20 slices (about 20 mm) with T2-weighted 3D cSSFSE. Right video shows various TI periods (250 to 3000 ms). Tagged CSF signals from the left side of brain outflow into the superior sagittal sinus (SSS), via along the lateral wall of the SSS.

